# Models trained to predict differential expression across plant organs identify distal and proximal regulatory regions

**DOI:** 10.1101/2024.06.04.597477

**Authors:** Michael C. Tross, Gavin Duggan, Nikee Shrestha, James C. Schnable

## Abstract

A large proportion of standing phenotypic variation is explained by genetic variation in noncoding regulatory regions. However, tools for the automated identification and characterization of noncoding regulatory sequences in genomes have lagged far behind those employed to annotate and predict the functions of protein coding sequences. We developed a modified transformer model and trained it to predict relative patterns of expression across a diverse set of tissues given a large sequence window for each gene of interest in the maize (*Zea mays*) genome. Nucleotides in the input DNA sequence with high saliency in gene expression pattern prediction overlapped with regions identified via comparative genomic or chromatin-based approaches as potential regulatory sequences. High saliency regions identified in a second species, sorghum (*Sorghum bicolor*), without species-specific training were also associated with potential regulatory sequences in noncoding regions upstream and downstream of each gene of interest. The potential impact of a scaleable and transferable approach to identifying regulatory sequences using saliency calculated from large context window models spans multiple applications. Specific use cases could include genome annotation, interpretation of natural genetic variation, and targeted editing in noncoding regions to alter patterns of levels of gene expression.

## Introduction

Gene editing has substantial potential to accelerate rates of genetic gain and engineer more resilient, resource-use-efficient, and nutritious crops(Nasti and Voytas 2021; Xu *et al*. 2017). However, achieving the full potential of gene editing will require fine-tuning levels and gene expression patterns and not only disrupting coding sequences, the most common editing approach used today in many crop and model species (Swinnen *et al*. 2016). The identification of protein-coding sequences within genomes and, to some extent, the prediction of which sequence changes will have greater or smaller impacts on protein function have become increasingly manageable tasks. However, identification of the noncoding regulatory regions that determine where, when, and in what quantities protein-coding DNA sequences are transcribed into mRNA, and the prediction of how changes in these regulatory sequences will influence function have remained far more challenging (Schmitz *et al*. 2022).

At least two distinct approaches to the identification of noncoding regulatory sequences have been used and validated on genome-wide scales: the identification of conserved noncoding sequences (Freeling *et al*. 2007), and the identification of open chromatin regions (Rodgers-Melnick *et al*. 2016). Comparison of orthologous genomic regions across related species can identify noncoding regions that, like many exons, exhibit slower rates of protein sequence evolution, indicating that these regions are likely to be functionally constrained. Several of these functionally constrained noncoding sequences (i.e. conserved noncoding sequences) have been shown to function in regulating the expression of nearby genes. However, by definition, evolutionarily young regulatory regions will not be identified as conserved noncoding sequences. In addition, the smallest sequences that can be confidently identified as showing functional constraints in plant genomes are substantially larger than known transcription factor binding sites. Conserved noncoding sequences thus mark a subset of the functional regulatory sequences present in a given plant genome. Open-chromatin regions identified via a range of sequencing-based methods – ATAC-seq, MNase-seq, etc – have been shown to contain a large majority of the genetic markers outside of gene bodies that are linked to variation in plant phenotypes (Rodgers-Melnick *et al*. 2016). However, current open-chromatin methods tend to identify larger genomic intervals and likely represent a superset of regulatory sequences in plant genomes.

Over the past five years, a range of increasingly complex machine learning algorithms have been employed for the task of predicting gene expression – defined in various ways – from DNA sequences. These models have demonstrated the ability to predict which of a pair of genes will be more expressed using DNA sequence starting one kilobase upstream of the annotated transcription start site and extending one kilobase downstream of the transcription end site in maize (Washburn *et al*. 2019) and even predict which genes will exhibit differential expression in response to external stress across maize and related species (Meng *et al*. 2021). Efforts to predict tissue-specific patterns of expression from DNA sequence have been less successful. A BERT-based transformer model exceeded controls but achieved prediction accuracies of R^2^=0.092-0.192 in predicting absolute expression levels in individual maize tissues (Levy *et al*. 2022). Another transformer-based effort was able to accurately estimate the overall strength of individual promoter elements but exhibited poor performance in predicting differences in expression of the same genes across different tissues from proximal promoter sequences (Mendoza-Revilla *et al*. 2023). Models that can predict gene expression patterns from DNA sequence can also be used to understand which DNA nucleotides play important roles in regulating gene expression and how DNA sequence changes may alter gene expression levels (Mendoza-Revilla *et al*. 2023).

## Results

We trained a transformer-based model to predict the relative expression of individual maize genes across six highly differentiated tissues using the sequence of the genomic interval starting 15 kilobases upstream of the gene’s transcription start site (TSS) and extending 15 kilobases downstream of the gene’s transcription end site (TES). We employed gene-family-guided-splitting (Washburn *et al*. 2019) to reduce data leakage between training and test data. To evaluate model performance we adopted the “Fortune Cookie Test” (see Extended Methods) as this control significantly outperformed more conventional controls, likely as a result of repeated patterns of tissue-specific expression across unrelated genes. Our trained model significantly outperformed the Fortune Cookie Test as assessed via average Spearman’s *ρ* between predicted and observed gene expression in 3,515 holdout test genes (7.349 x 10^−41^, Mann Whitney U test). Model predictive performance declined rapidly when smaller regions of noncoding sequence surrounding the gene of interest were employed for prediction and only exceeded that of controls when thirteen kilobases or more of surrounding noncoding sequence was provided to the model.

We evaluated the overlap between the saliency of individual DNA nucleotides when calculating relative tissue-level expression – estimated via a gradient-based approach – and two markers for DNA regulatory function: conserved noncoding sequences identified in comparisons between maize and rice (Turco *et al*. 2013) and open chromatin regions identified via ATAC-seq from leaf tissue (Lu *et al*. 2019). Comparisons were conducted using a set of 230 bigfoot genes that were associated with greater numbers of conserved noncoding sequences and typically exhibit more complex patterns of regulation and expression (Freeling *et al*. 2007). A number of maize-rice conserved noncoding sequences co-localized with spikes in saliency (Figure 1A), although not all spikes in saliency were tagged by conserved noncoding sequences nor did all conserved noncoding sequences correspond to spikes in the saliency map. The median base pair within an upstream or downstream conserved noncoding sequence exhibited significantly higher saliency than the median base upstream or downstream nucleotide, excluding conserved noncoding sequences, of the same gene (p=9.958 x 10^−7^, n=160 and p=4.625 x 10^−7^, n=163) respectively, paired t-test) (Figure 1B). Similarly, many but not all ATAC-seq peaks co-localized with spikes in the saliency maps for individual genes (Figure 1C) and the median base pair within ATAC-seq peaks exhibited significantly higher saliency than the median base pair in the remainder of the upstream or downstream regions of the same genes (p=1.545 x 10^−6^, n=106 and p=3.547 x 10^−5^, n=103) respectively, paired t-test) (Figure 1D). Unlike several recently described transformer-based models (Levy *et al*. 2022; Mendoza-Revilla *et al*. 2023), the model employed here was trained on data from only a single species, maize. Nevertheless, saliency scores calculated using the maize-trained model continued to be significantly associated with conserved noncoding sequence and open chromatin regions when the model was instead employed to predict relative patterns of tissue-level expression in a second related species, sorghum (Figure 1E-H). A general trend towards higher saliency scores downstream rather than upstream of the target gene is consistent with previous models (Washburn *et al*. 2019; Meng *et al*. 2021).

**Figure 1.**
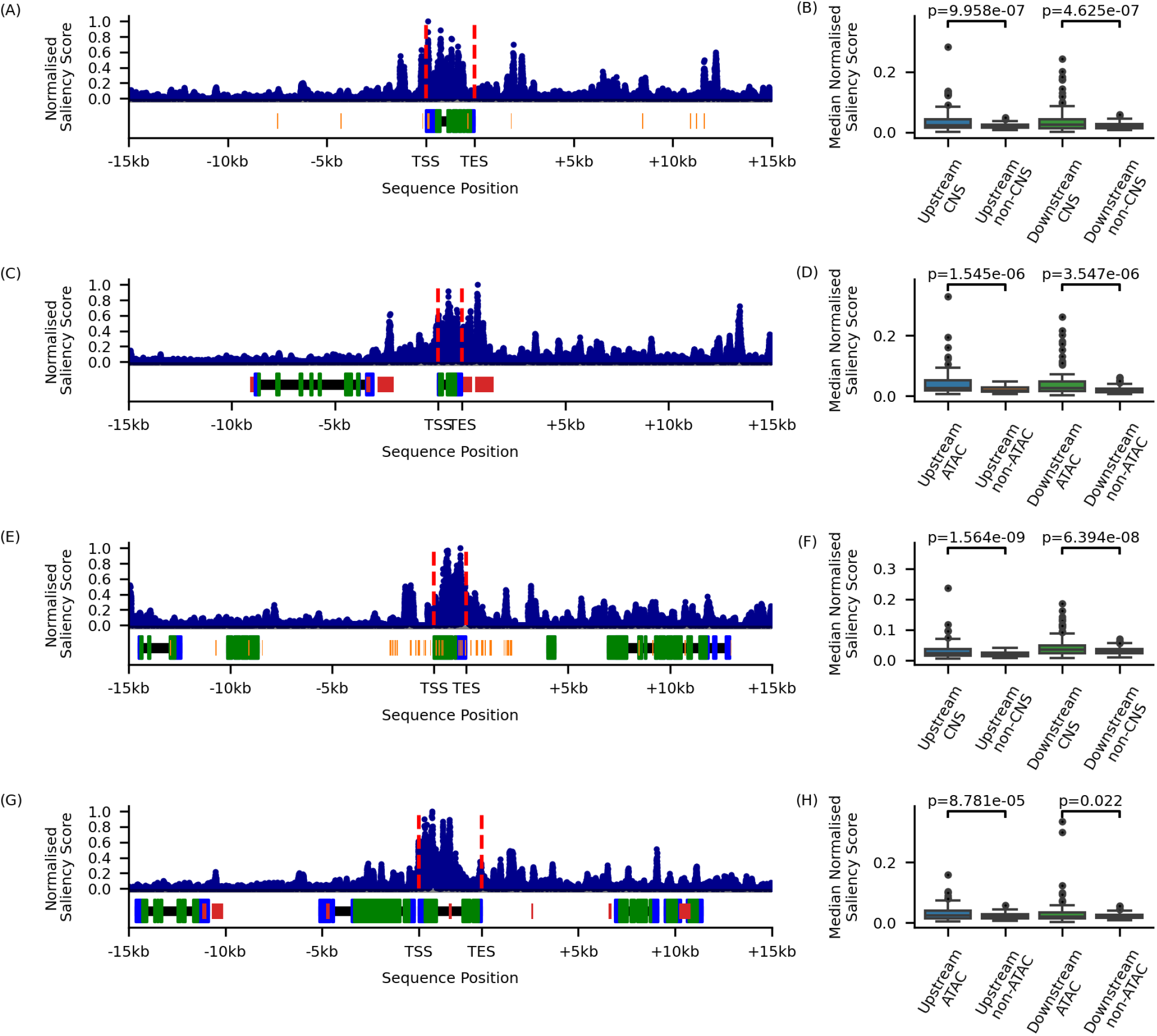
Association between saliency and marks of regulatory functions in maize and sorghum. A. Example of the distribution of maize-rice conserved noncoding sequences and saliency scores for a maize gene, GRMZM2G050933 encoding a Cyclin D6 protein in v2 of the maize reference genome. Black rectangles indicate the positions of annotated genes within the genomic interval used for prediction. Blue and green boxes indicate untranslated and translated exonic sequences, respectively. Blue dots indicate the max-normalized saliency score of the nucleotide at the corre-sponding position on the x-axis. Vertical dashed red lines indicate the annotated TSS and TES of the gene of interest. Orange lines indicate the positions of conserved noncoding sequences between maize and rice. B. Distribution of median max-normalized saliency scores for nucleotides located within and outside conserved noncoding sequences in the upstream (-15 kb to TSS) and downstream (TES to + 15 kb) regions of maize bigfoot genes with at least one annotated conserved noncoding sequence present in the respective regions. P-values shown were calculated via paired, two-tailed t-tests. Upstream region n = 160, downstream region n = 163. C. Example of the distribution of ATAC-seq peaks and saliency scores for a maize gene, Zm00001d006739 encoding a domain of unknown function (DUF260) protein in v4 of the maize reference genome. Red boxes indicate the position of annotated ATAC-seq peaks, all other features are as defined in panel A. D. Distribution of median max-normalized saliency scores for nucleotides located within and outside of the ATAC-seq peaks in the upstream (-15 kb to TSS) and downstream (TES to + 15 kb) regions of maize bigfoot genes with at least one ATAC-seq peak present in the respective region. P-values shown were calculated via paired, two-tailed t-tests. Upstream region n = 106, downstream region n = 103. E. Example of the distribution of sorghum-rice conserved noncoding sequences and saliency scores for a sorghum gene, Sb03g028390 encoding an uncharacterized protein in v1.4 of the sorghum reference genome. All features are as defined in panel A. F. Distribution of median max-normalized saliency scores for nucleotides located within and outside of the conserved noncoding sequences in the upstream (-15 kb to TSS) and downstream (TES to + 15 kb) regions of sorghum bigfoot genes with at least one annotated conserved noncoding sequence present in the respective regions. P-values shown were calculated via paired, two-tailed t-tests. Upstream region n = 185, downstream region n = 182. G. Example of the distribution of ATAC-seq peaks and saliency scores for a sorghum gene, Sobic.001G425300 encoding a lipase in v3 of the sorghum reference genome. All features are as defined in panel C. H. Distribution of median max-normalized saliency scores for nucleotides located within and outside of the ATAC-seq peaks in the upstream (-15 kb to TSS) and down-stream (TES to + 15 kb) regions of sorghum bigfoot genes with at least one ATAC-seq peak present in the respective regions. P-values shown were calculated via paired, two-tailed t-tests. Upstream region n = 114, downstream region n = 111. In panels B, D, F, and H color filled boxes represent the range between the 25th and 75th percentiles of median max-normalized saliency scores. Black solid lines inside colored boxes indicate the median (50th percentile) value. Whiskers extend to either the minimum and maximum values or 1.5 times the interquartile range. In the latter case, any data points beyond the edge of the whiskers are indicated with filled circles.

## Discussion

Machine learning-based approaches to predicting gene expression levels and patterns from associated DNA sequence are advancing rapidly. In many cases, only sequence from the proximal noncoding regions (e.g. one kilobase up and downstream of annotated exons) is employed for prediction, despite extensive evidence that more distal noncoding regions play key roles in determining gene expression pattern (Salvi *et al*. 2007; Studer *et al*. 2011). Here we employed sequence data from larger genomic intervals surrounding annotated genes to predict relative expression across a set of transcriptionally diverse tissues and demonstrate that per-nucleotide saliency scores extracted from this model are significantly associated with two different markers for functional noncoding regulatory sequence: the presence of conserved noncoding sequences and open chromatin. These results suggest that transformer-based architectures may provide a workable approach to guide efforts both to shape the transcriptional regulation of genes via gene editing beyond proximal promoter regions and understand the large proportion of standing phenotypic variation in populations attributable to genetic variances in noncoding regulatory sequence (Rodgers-Melnick *et al*. 2016).

## Materials AND Methods

An empirically derived custom transformer-based model described in extended methods was trained to predict the relative expression values of maize genes across six tissue types: leaf, root, anthers, immature cob, embryo, and endosperm. For each gene, expression data was normalized as a proportion of the maximum expression value across the six tissue types for that gene. Maize genes from the B73_RefGen_V5 reference genome were clustered into 10,738 unique gene families and whole gene families were assigned to training, testing, and validation datasets such that the total number of genes in each category approximated an 80%/10%/10% split. The input sequence for predicting each gene was the genomic interval beginning 15,000 base pairs upstream of the annotated transcription start site of the primary transcript and ending 15,000 base pairs downstream of the annotated transcription end site. Per nucleotide saliency in maize was calculated by taking the absolute gradient of the loss with respect to the input sequence where loss values were calculated via comparison of the predicted and observed relative expression profiles for the gene of interest. Expression and per nucleotide saliency values in sorghum were calculated similarly with the modification that loss was calculated relative to expression in six sorghum tissues identified as most similar to the six maize tissue-level expression datasets used in this study based on the degree of correlation in the absolute level of expression of syntenic orthologs between maize and sorghum expression datasets. Data on the positions of conserved noncoding sequences between the rice genome and syntenic orthologs in maize and sorghum were taken from (Turco *et al*. 2013). As these annotated conserved noncoding sequences were identified in earlier drafts of the maize and sorghum reference genomes, new saliency values were calculated using genomic intervals extracted from these earlier versions of the maize and sorghum reference genomes. Open chromatin windows defined via ATAC-seq were taken from leaf tissue in sorghum (Zhou *et al*. 2021) and leaf tissue in maize (Lu *et al*. 2019). More detailed information is available in Extended Methods.

## Code AND Data AVAILABILITY

The code used in this study is available at https://github.com/mtross2/transformer_regulatory_sequence.

## ACKNOWLEDGEMENTS

This work was supported by USDA-NIFA Awards No. 2020-68013-32371 & 2021-67021-35329 to JCS. This work was completed using the Holland Computing Center of the University of Nebraska, which receives support from the Nebraska Research Initiative.

## Author Contributions

MCT and JCS conceived of the project. MCT, JCS, NS generated, assembled, and quality controlled data. JCS and GD designed and advised on the analysis methods. MCT and JCS conducted the analyzes. MCT and JCS wrote the first draft of the manuscript. GD and NS contributed significant additional content during the revision of the manuscript. All authors read and approved the final version of the manuscript.

## Author Declaration

Michael C. Tross and James C. Schnable are listed as inventors on U.S. Patent Application 63654,690, “A computational method for identifying specific DNA sequences that regulate the level and pattern of RNA expression for genes of interest”. The authors declare no other conflicts of interest.

## Supplementary Information

### Extended Methods

#### Extracting and processing of genomic sequence intervals

Both genomic sequence and GFF3 annotation files for multiple maize reference genome versions (B73_RefGen_V2, B73_RefGen_V4, and B73_RefGen_V5) (Schnable *et al*. 2009; Jiao *et al*. 2017; Hufford *et al*. 2021) and the sorghum reference genome version (BTx623 v3.1) (McCormick *et al*. 2018) were downloaded from Phytozome 13 (Goodstein *et al*. 2012). Genomic sequence and GFF annotations for the BTx623 v1.4 version of the sorghum reference genome (Paterson *et al*. 2009) were downloaded from CoGE comparative (Lyons *et al*. 2008).

For each gene of interest in each genome version, the region beginning 15,000 base pairs upstream of the annotated start position of the gene and extending 15,000 base pairs downstream of the annotated end position of the gene was extracted. For genes on the reverse strand, the extracted sequence was reverse complemented. When multiple alternative splice variants were annotated for the same gene, the earliest annotated transcription start site and the last annotated transcription end site were used to define the start and the end of the gene for purposes of extracting sequence. Genes for which the criteria above resulting in sequences >90,000 base pairs were excluded from training, validation and testing.

#### Source and processing of gene expression datasets

Data on the expression of annotated maize genes across 57 distinct RNA-seq datasets (Stelpflug *et al*. 2016) were sourced from MaizeGDB (Portwood *et al*. 2019). Data on the expression of annotated sorghum genes across 49 distinct RNA-seq datasets generated as part of seven previously published studies (Emms *et al*. 2016; Kebrom *et al*. 2017; Makita *et al*. 2015; Davidson *et al*. 2012; Olson *et al*. 2014; Turco *et al*. 2017; Wang *et al*. 2018), reanalyzed and re-quantified using the iRAP pipeline, were sourced from the EBI Expression Atlas (Moreno *et al*. 2022).

The six pairs of a maize expression dataset and a sorghum expression dataset were selected using a combination of empirical knowledge about functionally distinct tissue types and an analysis of the degree of correlation in the absolute expression of 11,108 maize-sorghum syntenic orthologous gene pairs to identify pairs of tissue with similar patterns of gene regulation between the two species (Zhang *et al*. 2017). The six final pairs of expression data and the original sample names using in their respective sources where:

- **Leaf**: ‘V7 Tip of transition leaf maize’ (maize), ‘flag leaf’ (sorghum)
- **Root**: ‘Primary Root 6 days after sowing GH’ (maize), ‘seedling developmental stage, root’ (sorghum)
- **Inflorescence**: ‘V18 Immature cob’ (maize), ‘inflorescence size 1 to 10 millimeter, inflorescence’ (sorghum)
- **Anther**: ‘Anthers R1’ (maize), ‘anther’ (sorghum)
- **Embryo**: ‘Embryo 22 DAP’ (maize), ‘plant embryo’ (sorghum)
- **Endosperm**: ‘Endosperm 24 DAP’ (maize), ‘20 days after pollination, endosperm’ (sorghum)

#### Family guided gene splitting for datasets

Maize genes were divided into training, validation, and testing datasets using gene family guided splitting (Washburn *et al*. 2019) as described below. Maize gene family assignments, including single gene families and families including as many as 352 gene copies were extracted from PLAZA (Van Bel *et al*. 2022). The set of 10,738 unique maize gene families containing at least one expressed maize gene were subdivided into training, validation, and testing data sets with a target of 80%, 10%, 10% of individual genes present in each category. The final split contained 28,076, 3,521 and 3,515 genes in the training, validation and test sets respectively, with no single gene family represented in two or more of the three sets.

### Architecture and training of transformer-model

An empirically derived custom transformer-based model with input convolutional layers was implemented in python using Pytorch (V2.1.0) software (Paszke *et al*. 2017). There was six convolutional layers, all having kernel sizes of 15, and 1000, 500, 250, 500, 500, 1000 feature maps respectively. The 2nd, 4th and 6th convolutional layers were each followed by RELU activation function, a max pooling layer of a kernel size of 2 and step size of 2, then a dropout layer of 0.25 before the succeeding convolutional layers. All other convolutional layers was followed only by a RELU activation function. The output of the final convolutional layer was fed into a positional encoding layer then five multi-attention layers each having 1000 embedding dimensions, 8 attention heads, and a skip connection incorporating information from the positional encoding layer. Layer normalization was performed on each multi-head attention layer. Finally the output of the series of multi-attention heads were fed into three fully connected layers of sizes 4000, 1000, and 6 respectively. A Binary Cross Entropy loss function which took logits as input and applied a sigmoid activation function, was used to calculate the loss between the predicted and observed expression profile of each gene. An Adam optimizer was used to optimize the weights of the model towards a lower loss, using a starting learning rate of 0.000001.

For each gene in the training dataset, the genomic sequence extracted above was one-hot encoded based on four possible nucleotide sequences (A, T, C, and G). X or N values indicative of gaps or masked sequence in the input sequence were represented as all 0s across the four variables encoding a given nucleotide position. The expression profile for each gene in the training, validation, and testing datasets used for loss calculations was generated by dividing the expression of each gene in a given RNA-seq dataset by the maximum observed expression for that same gene across all RNA-seq datasets employed.

For training, backpropagation was performed on a gene by gene basis. The model was trained for 207 epochs, at which point no further improvement in model performance was observed after an additional 100 epochs. The final model used was the best performing one as assessed on hold out validation data (described above).

### Controls for model predictive performance

The performance of the trained transformer model in predicting patterns of gene expression across tissues was quantified via Spearman’s *ρ* between predicted and observed expression profiles on the 3,515 maize genes of the hold out test dataset. The observed distribution of *ρ* values were compared to three different controls, with control three (the “Fortune Cookie Test”) exhibiting the highest baseline performance.

- Control 1: *ρ* calculated between the predicted expression for a given gene and the shuffled versions of the true set of expression values for the same gene.
- Control 2: *ρ* calculated using the observed set of expression values for the gene of interest and the observed set of prediction values for another, randomly selected gene, as a stand in for the predicted expression values.
- Control 3 (“Fortune Cookie Test”): *ρ* calculated using the observed set of expression values for the gene of interest and a set of predicted expression values produced by the model for another, randomly selected gene.

#### Saliency maps

Sequence for gene of interest was extracted and pre-processed as described for model training. The trained transformer model was employed to predict relative expression values using the sequence for the gene of interest as input. The loss value as determined by the model loss function, between the predicted and observed relative expression values was computed. The gradient of the loss with respect to the input sequence was derived for each input nucleotide sequence – the absolute average of the four input values used to represent each nucleotide via one-hot encoding – and is referred to below as the saliency score. Ambiguous nucleotides (X and N) was masked for downstream analyses. Normalized (relative) saliency scores for individual genes were determined by max normalization of all nucleotide-level scores for the gene of interest.

### Ground truth gene regulatory sequence data sources

Conserved noncoding sequences used in this study were sourced from a previous study and identified via comparison of the genomic intervals starting 12,000 bp upstream of individual syntenic orthologous gene pairs between either maize (B73_RefGen_V2) or sorghum (BTx623 1.4) and rice (MSU v6) and extending 12,000 bp downstream of the same individual gene pairs (Turco *et al*. 2013).

The maize leaf ATAC-seq peak dataset used in this study was originally described in (Lu *et al*. 2019) and downloaded via the Plant Epigenome Browser. The sorghum leaf ATAC-seq peak dataset used in this study was originally described in (Zhou *et al*. 2021) and downloaded from the publisher’s website as “Supplemental Table 2: ACRs in sorghum genome.” Comparisons of the saliency scores of putative regulatory sequences – CNS or ATAC-seq peaks – with control sequences were performed for a set of 230 maize bigfoot genes – defined here as genes having more than 10 CNS regions within the 12,000 bp upstream and 12,000 bp downstream interval of the genebody – and a set of 332 sorghum bigfoot genes – genes having more than 15 CNS regions within the 12,000 bp upstream and downstream interval of the genebody –.

The median of normalized saliency scores within putative regulatory sequences was determined for all big foot genes. Saliency was partitioned into scores for nucleotides that overlapped conserved noncoding sequence or ATAC-seq peaks and nucleotides which did not overlap. Upstream and downstream regions were considered independently and median saliency scores for overlapping and nonoverlapping nucleotides were only calculated for a given region if at least one putative regulatory sequence – CNS or ATAC-seq was present in that region for that gene.

